# A General Framework for Injecting Biophysical Priors into Protein Embeddings

**DOI:** 10.64898/2025.12.23.696257

**Authors:** Jonathan Feldman, Antoine Maechler, Dianzhuo Wang, Eugene Shakhnovich

## Abstract

Accurate ΔΔ*G* prediction requires integrating machine learning with biophysical insight. Existing approaches typically prioritize one while neglecting the other. We introduce an encoder-agnostic framework that injects interpretable biophysical priors into residue-level deep learning representations via cross-embedding attention. ProtBFF consistently improves performance under homology-based-clustering evaluation and enables general-purpose encoders to surpass state-of-the-art specialized models or larger models. Our results show that integrating simple, mechanistic priors into pretrained representations yields more trustworthy predictors, offering a practical solution for broader protein engineering applications.

**Code:** github.com/Jfeldman34/ProtBFF

## Introduction

Predicting how multiple mutations alter protein–protein binding affinity (ΔΔ*G*) is a central challenge in computational biology with direct implications for protein engineering Sanavia et al. (2020); Pucci et al. (2022); Huot et al. (2025); Wang et al. (2024). Biophysics-based methods such as molecular dynamics have long been used for this task, but they are computationally expensive and rely on hard-coded mathematical models, which limits their scalability Schymkowitz et al. (2005); Park et al. (2016); Alford et al. (2017). While deep learning has transformed other realms of protein science, such as protein structure prediction Abramson et al. (2024); Hayes et al. (2024); Watson et al. (2023); Feldman and Skolnick (2025), the prediction of ΔΔ*G* remains constrained by small, biased datasets and by models that often fail to generalize to previously unseen proteins Hummer et al. (2025); Loux et al. (2024). This shortcoming arises from two interconnected factors. First, the quality and quantity of available experimental measurements are limited, and existing datasets cover only a narrow diversity of proteins. Second, many models tend to overfit dataset-specific patterns rather than learning the underlying biophysical principles that govern protein–protein interactions.

A critical example is the SKEMPI2 dataset, the field’s most used benchmark. While SKEMPI2 contains roughly 350 curated protein complexes, it suffers from substantial sequential and structural redundancy, with many complexes exhibiting high structural and sequential similarity Bushuiev et al. (2024); Jankauskaitė et al. (2018). As this work will show, such hidden redundancy introduces significant data leakage between training and test sets, inflating reported performance and obscuring the true generalization gap in current methods Bushuiev et al. (2024); Jankauskaitė et al. (2018).

Although it is widely appreciated that biophysical insight should guide protein binding prediction, there remains no clear, general strategy for injecting such priors into modern deep learning models. To this end, we introduce ProtBFF (Protein Biophysical Feature Framework), an encoder-agnostic module that enriches embedding-based predictors with explicit biophysical features. Unlike prior work that proposes entirely sophisticated architectures Luo et al. (2023); Yu et al. (2024); Mo et al. (2024) or relies solely on physics-based scoring functions Schymkowitz et al. (2005); Al-ford et al. (2017), ProtBFF reweights residue-level embeddings based on their local structural and physico-chemical context. (Figure 1). By injecting biophysical priors directly into the latent space, ProtBFF biases learning toward features that are informed by known physical determinants of protein–protein interactions while retaining the flexibility of data-driven models. ProtBFF acts as a plug-in that integrates seamlessly with any pretrained encoder. We demonstrate that it improves performance on SKEMPI2 and out-of-distribution antibody-antigen datasets, enables models not originally designed for this task, such as ProSST Li et al. (2024b) and the ESM family Hayes et al. (2024), to reach or surpass state-of-the-art specialized PPI predictors.

**Figure 1.**
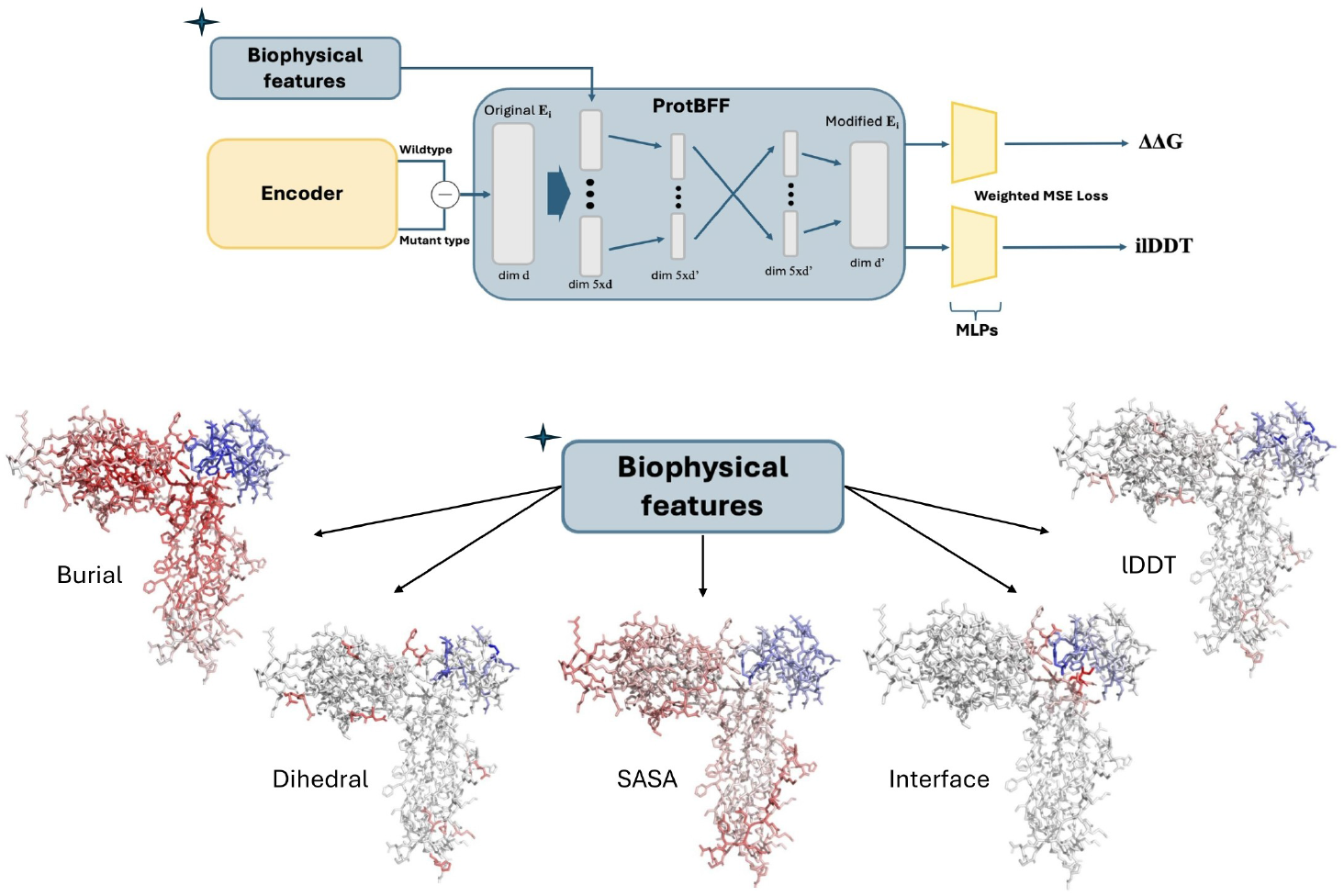
The upper part of the figure shows a simplified schematic of the ProtBFF framework. The pipeline begins with embedding extraction, followed by embedding subtraction and scaling using biophysical features. These processed embeddings are then passed through a pooling layer, a cross-embedding attention module, and finally two MLP heads to generate predictions of ΔΔ*G* and ilDDT, optimized via a weighted loss function. The lower part of the figure illustrates the five biophysical features by coloring residues according to their scores, with chain A shown in red and chain B in blue. Residues with higher biophysical contributions are colored more intensely, while those with weaker contributions appear closer to gray.

## Limitations of the SKEMPI2 Dataset

### Data Leakage

The main issue with the SKEMPI2 dataset is systematic data leakage. Prior studies mostly split train and test sets simply by complex PDB ID Luo et al. (2023); Yu et al. (2024); Mo et al. (2024). However, many differently named complexes are highly homologous in sequence and structure Bushuiev et al. (2024); Jankauskaitė et al. (2018); Yu et al. (2025) (Figure 2 B). As a result, homologous proteins often appear in both splits, inflating benchmark performance.

**Figure 2.**
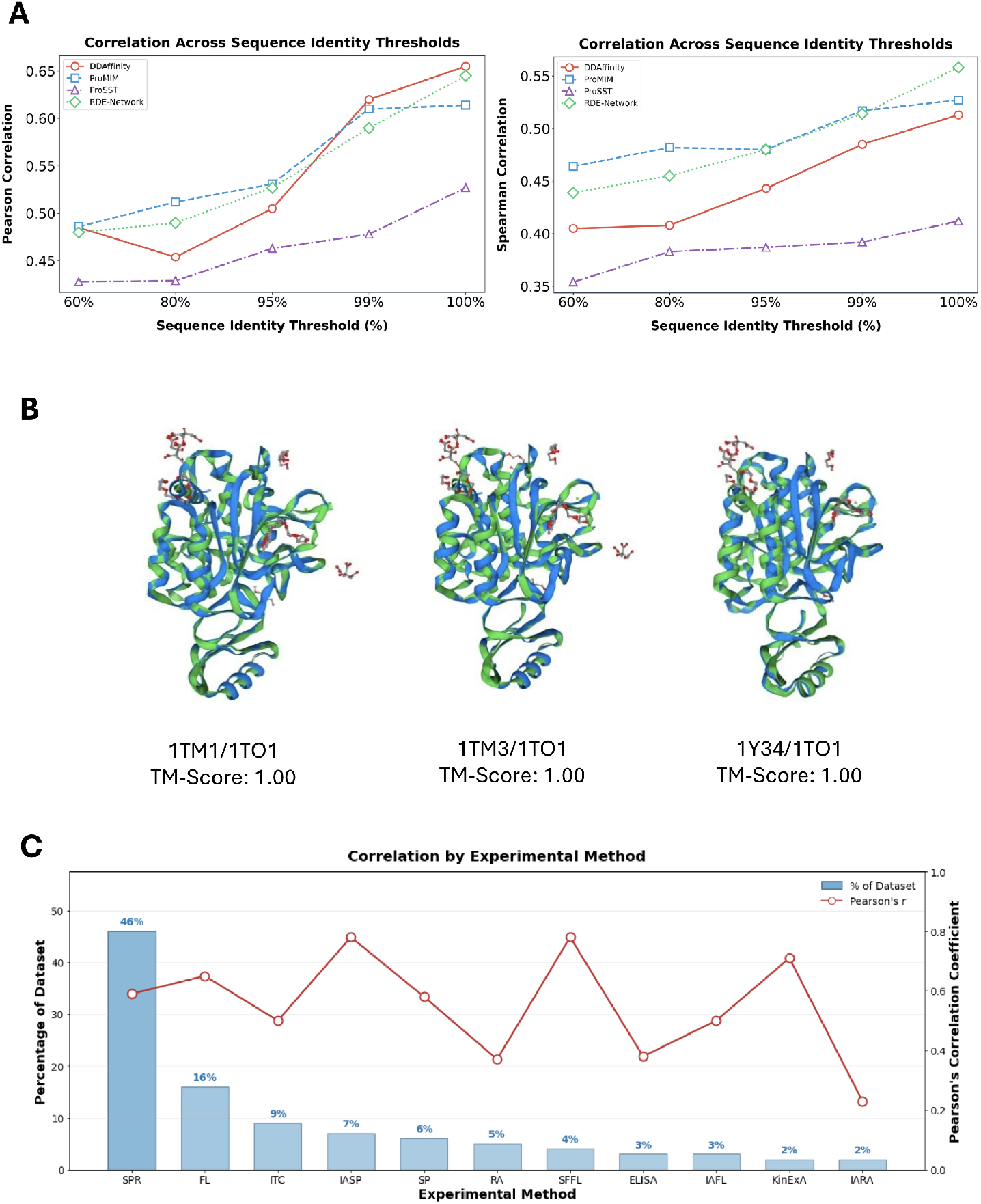
**A** Performance of deep learning ΔΔ*G* predictors on SKEMPI2 under progressively stricter sequence-identity clustering. Lines report correlation for each model, with the left subfigure being the Pearson correlation and the right being the Spearman correlation. Model performance declines sharply as homology constraints are enforced. **B** Structural superposition of three representative SKEMPI2 protein complexes (1TM1, 1TM3, and 1Y34) onto a fourth complex using US-Align Zhang et al. (2022). Although treated as distinct dataset entries, these complexes share over 99% sequence identity and exhibit near-identical structures, illustrating the extent of hidden redundancy in SKEMPI2. **C** Leave-one–experimental-method-out evaluation of the ΔΔ*G* predictor DDAffinity on SKEMPI2. For each experimental method, the model was trained on all datapoints except those measured using the held-out method and tested exclusively on that method. Only experimental methods comprising at least 1% of the dataset are shown. To isolate effects attributable to experimental methodology, homology-based clustering was not applied in this analysis. Pearson’s correlation coefficient is shown on the y-axis, with experimental methods on the x-axis.

To quantify this effect, we clustered SKEMPI2 complexes by sequence identity (see Methods), applying thresholds from 60% to 100%. Complexes whose chain sequences exceeded a given threshold were grouped together; at 100% identity this is equivalent to no clustering. As the threshold is lowered, the number of clusters decreases rapidly, revealing extensive homology within the dataset. Notably, reducing the threshold from 100% to 99% reduces the number of clusters from 335 to 253, and at 60% identity only 136 clusters remain (Table 1), demonstrating substantial redundancy that is not removed by splitting complexes by name alone.

**Table 1.** Number of protein complex clusters at different sequence identity thresholds. Lower thresholds merge homologous complexes, highlighting the redundancy in SKEMPI2.

| Threshold | 60% | 80% | 95% | 99% | 100% |
| --- | --- | --- | --- | --- | --- |
| Clusters | 136 | 179 | 219 | 253 | 335 |

Beyond sequence similarity, SKEMPI2 aggregates ΔΔ*G* measurements obtained using diverse experimental techniques with varying levels of reliability Jankauskaitė et al. (2018), which introduces additional noise into the dataset. As shown in Figure 2 C, model performance depends strongly on the experimental method used for measurements. In particular, indirect or surface-based assays, such as Enzyme-Linked Immunosorbent Assay (ELISA), can introduce substantial measurement noise **?**. This increased experimental variance degrades predictive accuracy and makes the corresponding data more difficult for models to learn from.

### Impact of Data Leakage on Models’ Performance

To quantify the effect of hidden leakage, we bench-marked four common models (DDAffinity Yu et al. (2024), ProMIM Mo et al. (2024), RDE-Network Luo et al. (2023), ProSST Li et al. (2024b)) using 10-fold cross-validation on sequence-clustered SKEMPI2. As the sequence-identity threshold is lowered, train–test homology falls and all models’ Pearson and Spearman correlations drop markedly (Fig. 2A). This trend highlights that previously reported results on SKEMPI2 were strongly influenced by redundancy between training and test sets, ultimately leading to overfitting.

## Systematic Way of Injecting Biophysical Priors

### Prediction Improvements with ProtBFF

To assess the improvements provided by ProtBFF as a plug-in module, we integrated it into two types of models that follow the latent representation paradigm and benchmarked them against several established baselines: ProSST, originally developed for single-protein stability prediction Li et al. (2024b), and the ESM2 Lin et al. (2023) and ESM3 models Hayes et al. (2024), two general-purpose protein language models (the first one sequence-only, and the other integrating sequences and structures). All models were retrained using a 10-fold cross-validation protocol on the SKEMPI2 dataset clustered at 60% sequence identity, as detailed in the Methods section

ProSST Li et al. (2024b), though not originally designed for protein complex ΔΔ*G* prediction, improves substantially with ProtBFF, with the Pearson increasing from 0.428 to 0.515 and the Spearman increasing from 0.354 to 0.471, surpassing specialized models such as ProMIM and DDAffinity. ESM models also benefit substantially, with Pearson and Spearman correlations increasing to levels comparable to most state-of-the-art models.

These results illustrate that models pretrained on large protein datasets, such as the CATH database Knudsen and Wiuf (2010) for ProSST and the extensive sequence and structural corpora underlying ESM2 and ESM3, learn rich general-purpose representations Hayes et al. (2024); Li et al. (2024b). By guiding these embeddings with ProtBFF, which selectively amplifies structurally relevant residues through biophysical features, we are able to unlock performance that rivals or even surpasses methods specifically designed for protein complex ΔΔ*G* prediction. Further information about the benchmarking and integration process can be found in the Methods section.

We also evaluated the effect of applying ProtBFF across the ESM2 model family, ranging from 150 million to 15 billion parameters. With ProtBFF, the 150-million-parameter model notably outperforms even much larger counterparts, as well as all other standard ESM2 variants. ProtBFF also provides consistent performance gains across all model sizes (Figure 3). Interestingly, the standard ESM2 650M model outperforms the 3B and 15B variants on SKEMPI2, consistent with observations that smaller ESM models sometimes match or exceed larger ones on specific predictive tasks Vieira et al. (2025).

**Figure 3.**
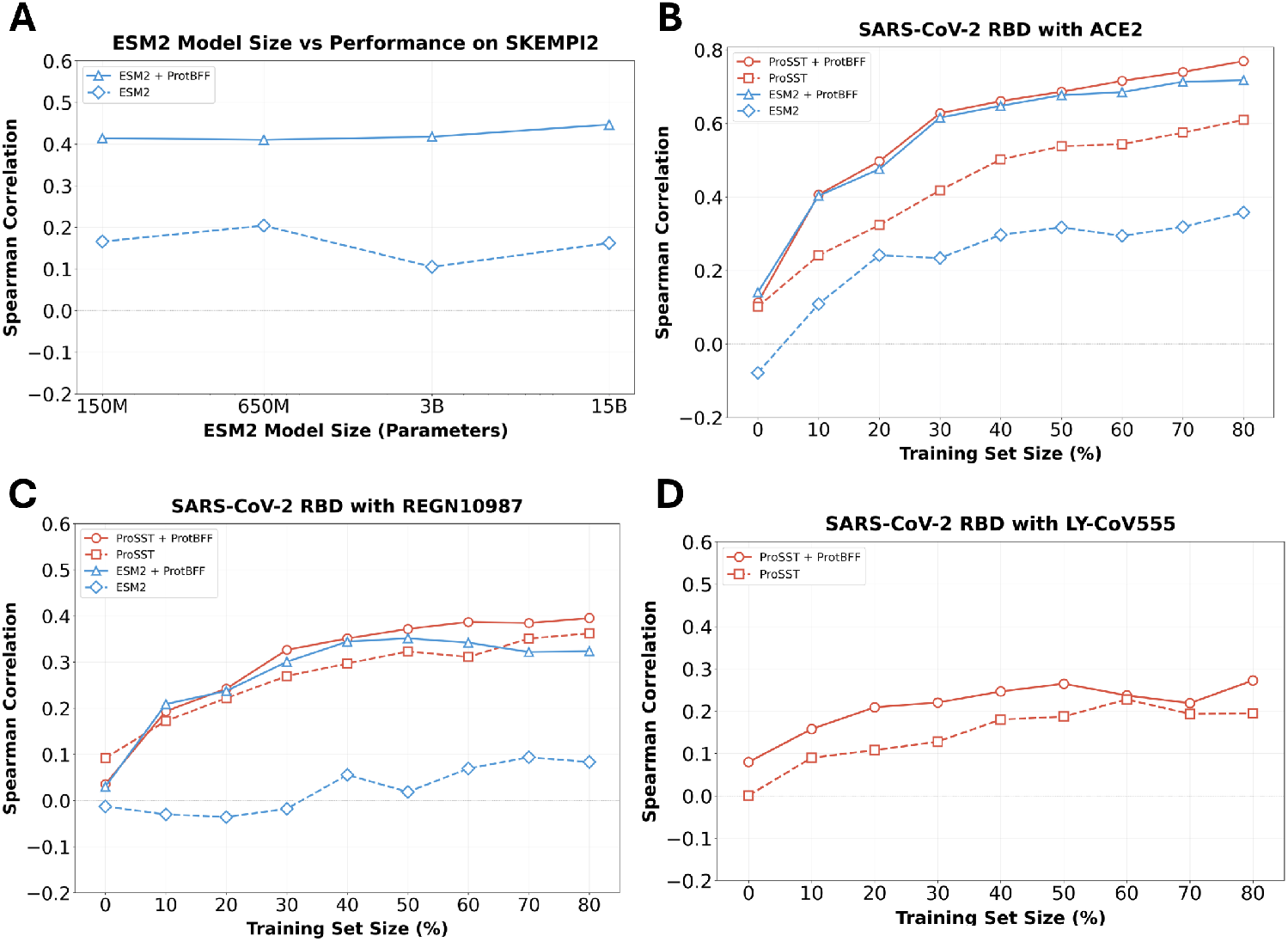
**A** Performance of ESM2 on the SKEMPI2 dataset, measured by Spearman correlation coefficient across model sizes defined by the number of parameters. **B–D** Performance of ProSST and ESM2, measured by Spearman correlation coefficient, with and without ProtBFF, on deep mutational scanning (DMS) datasets of SARS-CoV-2 RBD mutations when bound to **B** ACE2, **C** REGN10987, and **D** LY-CoV555. *Note:* Due to the large size of the RBD–LY-CoV555 complex template structure (PDB 7KMG), it exceeds the context length limit of ESM2 and therefore could not be processed. As a result, ESM2-based results are omitted from subfigure D. Refer to Supplementary Figure S1 for the Pearson correlation coefficients for these analyses.

#### Ablation Study Results

To assess the contribution of the five biophysical scores and the auxiliary ilDDT loss to ProtBFF’s performance, we conducted an ablation study. Each component was systematically removed in turn, and we observed that all scores as well as the ilDDT loss provided measurable improvements in both Pearson and Spearman correlations. Among them, the interface and burial features contributed the largest gains. These findings demonstrate that ProtBFF’s accuracy arises from the integration of multiple complementary biophysical signals rather than reliance on any single feature.

Table 2 presents the effects of systematically removing each biophysical feature from ProtBFF. All features contribute positively to predictive performance, with the interface and residue scores showing the largest impact; removing these two features leads to substantial drops in both Pearson and Spearman correlations, confirming their central role in capturing mutational effects on binding affinity. The dihedral, SASA, and lDDT scores produce smaller, yet still meaningful, reductions in performance, indicating that they provide complementary structural information that refines the model’s predictions. Additionally, removing the ilDDT-based loss function results in a measurable decline in both correlation metrics, demonstrating that incorporating structural fidelity into the training objective improves model robustness. Finally, baseline variants without any biophysical scores or embeddings (i.e. with all biophysical scores or embeddings set to 1) perform significantly worse, highlighting that the multi-feature integration strategy is crucial for ProtBFF’s predictive power.

**Table 2.** Benchmarking and ablation results. Top: Performance of baseline encoders with and without ProtBFF. Bottom: Ablation study of ProtBFF applied to ProSST, where each row removes one feature or the auxiliary *ilDDT* loss. Values reported are Pearson (*ρ*) and Spearman (*r*) correlations. Bold indicates best performance. Asterisks (*) mark methods where ProtBFF is not applicable.

| Benchmarking and Ablation Results<br>Method / Variant | Baseline |  | With ProtBFF |  |
| --- | --- | --- | --- | --- |
| | Pearson ($\rho$) | Spearman ($r$) | Pearson ($\rho$) | Spearman ($r$) |
| *Model Benchmarking* |  |  |  |  |
| ProSST Li et al. (2024b) | 0.428 | 0.354 | **0.515** | **0.471** |
| ESM2 Lin et al. (2023) | 0.194 | 0.204 | 0.451 | 0.410 |
| ESM3 Hayes et al. (2024) | 0.159 | 0.099 | 0.362 | 0.347 |
| ProMIM Mo et al. (2024) | 0.486 | 0.464 | * | * |
| RDE-Network Luo et al. (2023) | 0.480 | 0.439 | * | * |
| DDAffinity Yu et al. (2024) | 0.485 | 0.405 | * | * |
| RDE-Linear Luo et al. (2023) | 0.369 | 0.360 | * | * |
| FoldX Schymkowitz et al. (2005) | 0.320 | 0.294 | * | * |
| *Ablation Study on ProSST* |  |  |  |  |
| None (Full ProSST+ProtBFF) | — | — | **0.515** | **0.471** |
| Interface Score removed | — | — | 0.462 | 0.418 |
| Burial Score removed | — | — | 0.471 | 0.426 |
| Dihedral Score removed | — | — | 0.498 | 0.458 |
| SASA Score removed | — | — | 0.503 | 0.464 |
| IDDT Score removed | — | — | 0.506 | 0.469 |
| iIDDT Loss Function removed | — | — | 0.505 | 0.459 |
| **All Biophysical Scores set to 1** | — | — | 0.436 | 0.385 |
| **Protein Embeddings set to 1** | — | — | 0.077 | 0.123 |

#### SARS-CoV-2 Dataset Results

We next tested how well ProtBFF-enhanced models generalize to unseen tasks, focusing on ΔΔ*G* prediction for virus-receptor and antibody–antigen binding, which are key to antibody design and pandemic preparedness Feldman and Feldman (2025); Wang et al.; Yin and Pierce (2024). Performance was evaluated on a DMS dataset measuring SARS-CoV-2 RBD binding to ACE2, LY-CoV555, and REGN10987 Starr et al. (2020); Fowler and Fields (2014). Pretrained ProSST- and ESM2-based models, with and without ProtBFF, were assessed under zero-shot and few-shot learning across varying training set sizes, as shown in Figures 3 A–C.

This expanded benchmarking yields two key insights. First, ProtBFF consistently improves both ProSST and ESM2 representations when paired with a simple classifier. Across datasets, ProSST with ProtBFF often outperforms its base version by over 0.2 in Pearson and Spearman correlations, while ESM2 gains even more, approaching the performance of ProSST with ProtBFF. Second, although all models perform poorly in the zero-shot setting because SKEMPI2 provides limited coverage of antibody–antigen and virus–receptor interactions, few-shot learning markedly improves accuracy. With only 10% of training data, ProtBFF-enhanced models achieve high predictive performance, suggesting practical utility in data-limited contexts such as active learning.

Notably, the strong fine-tuned performance reflects the dataset’s structure. Evaluations are on DMS data from a single protein complex, making prediction easier than cross-complex generalization. Repeated exposure to mutations on the same backbone allows the models to leverage ProtBFF’s biophysical features more effectively, enhancing embeddings and boosting performance metrics.

## Discussion

Predicting mutational effects on protein–protein binding exposes a fundamental challenge for machine learning in biology: reliable generalization from small, biased datasets. Despite major advances from large-scale self-supervised pretraining Abramson et al. (2024); Hayes et al. (2024); Watson et al. (2023), supervised binding ΔΔ*G* prediction remains data-limited, with benchmarks such as SKEMPI2 comprising only 7,085 measurements across 345 complexes Pucci et al. (2022); Bushuiev et al. (2024); Jankauskaitė et al. (2018); Deng et al. (2025). Our analysis shows that these limitations are compounded by pervasive sequence and structural redundancy, which leads to homology-driven data leakage and inflated performance estimates. When evaluated under sequence-identity–based clustering, predictive accuracy drops sharply across methods, revealing that many models rely in part on memorization rather than transferable biophysical understanding.These results highlight the need for benchmarks and models that emphasize mechanistic understanding. Pretrained embeddings alone usually are insufficient in low-data regimes and can be enhanced by injecting physically interpretable, task-relevant priors.

ProtBFF provides a simple framework for injecting arbitrary biophysical priors into pretrained protein embeddings without discarding residue-level information through premature pooling. Across encoder architectures, ProtBFF consistently improves performance by integrating biophysical signals directly into the latent space. Ablation experiments show that these gains arise from the combination of pretrained embeddings and complementary biophysical priors, rather than from any single component alone. For the task of predicting binding affinity, interface and burial descriptors contribute the most, while global structural features such as solvent accessibility, dihedral geometry, and lDDT provide smaller but consistent improvements by adding complementary context.

ProtBFF should be viewed as an initial step rather than a complete solution. The five biophysical features used here provide only a coarse approximation of the physical processes governing binding, and richer sources of physical information could be incorporated within the same framework. Importantly, ProtBFF places no inherent limit on the number or type of biophysical priors: additional energetic, structural, or dynamic descriptors can be integrated, with the model learning how to weight and combine them from training data **?**. In this work we focus on protein–protein binding ΔΔ*G*, but the framework is general and could be readily extended to related problems such as protein folding stability, ligand binding, or fitness prediction **??**. Fully capturing dynamical effects beyond the fixed-backbone approximations of FoldX Schymkowitz et al. (2005); Alford et al. (2017) remains an important direction for future work.

ProtBFF operates on encoder models that produce residue-level embeddings, which makes it natural to incorporate biophysical information as a modular add-on aligned to individual residues. In contrast, more complex architectures that rely on higher-order geometric or relational representations may not admit such a straight-forward augmentation, and may require physical priors to be integrated directly into the model design Yu et al. (2024); Marquet et al. (2022). This distinction highlights a broader challenge in protein machine learning: determining when physical knowledge can be injected as a modular constraint, and when it must be co-designed with the representation itself.

Beyond model design, our findings underscore the importance of rigorous dataset construction and evaluation. Without careful control for redundancy, increasingly sophisticated models risk becoming effective memorization systems with limited practical relevance Graber et al. (2025). Developing larger, more diverse, and experimentally robust datasets is therefore critical for translating methodological advances into reliable predictive tools.

## Materials and Methods

### Dataset Collection

#### SKEMPI Dataset Preprocessing

In this study, we used the SKEMPI2 dataset for both benchmarking and training. SKEMPI2 contains 345 protein complexes and 7,085 combinatorial point mutations with experimentally determined ΔΔ*G* values, with structures obtained from the Protein Data Bank (PDB) Jankauskaitė et al. (2018). Due to input size limits of the smallest benchmarked model, ProSST (which was not trained to encode protein complexes), the 10 largest complexes were excluded Li et al. (2024b). This yielded a final working set of 335 complexes and 6,631 mutational variants.

To ensure fair comparisons, this reduced SKEMPI2 dataset was used consistently for all training and bench-marking. We applied ten-fold cross-validation using identical splits across all experiments. Before generating the splits, we clustered sequences with CD-HIT to ensure that complexes in different folds shared less than a specified sequence homology threshold Li and Godzik (2006). This step mitigated the data leakage issue in SKEMPI, since high sequence similarity often implies structural similarityKrissinel (2007); Koehl and Levitt (2002). Given that SKEMPI2 contains many complexes with nearly identical composition, this clustering also reduced structural homology overlap across folds Jankauskaitė et al. (2018).

### ProtBFF architecture

A central paradigm in molecular biology is to learn expressive latent representations of molecules and their interactions, while using relatively simple models for downstream prediction tasks such as ΔΔ*G* Marquet et al. (2022); Li et al. (2024a); Dahal et al. (2025). Concretely, an encoder generates per-residue embeddings for both wildtype and mutant proteins, integrating sequential and or structural information Mo et al. (2024); Luo et al. (2023); Yu et al. (2024). These encoders are typically pretrained on large-scale datasets with self-supervised objectives (e.g., masked residue prediction) to build informative internal representations Marquet et al. (2022); Li et al. (2024b). In this study, we propose a general biophysical framework that augments such models by integrating explicit biophysical features into per-residue embeddings, which can be seen in Figure 1.

Our framework builds on the observation that embedding-based models produce a latent representation for every residue, yet not all residues contribute equally to changes in binding free energy. Prior studies have shown that simple per-residue biophysical scores can strongly correlate with mutational effects on binding affinity Yu et al. (2024); Dauparas et al. (2022). Motivated by this, we enrich residue embeddings by scaling them with biophysical metrics: embeddings of residues with high structural or interfacial relevance are amplified, while those less likely to contribute are down-weighted. Incorporating multiple complementary scores enables the model to integrate diverse structural cues, while cross-embedding attention allows these signals to interact and emphasize the most informative patterns Girdhar and Ramanan (2017); Tang et al. (2023). Together, this yields richer and more interpretable residue-level representations. To implement this idea, we generate five scaled copies of each residue embedding, each weighted by a distinct biophysical metric: interface propensity, residue burial, dihedral deviation, solvent-accessible surface area (SASA), and local distance difference test (lDDT), which are visualized in Figure 1. These metrics are computed from the wildtype structures and FoldX-generated mutant structures Schymkowitz et al. (2005). Formally, for residue *i* with embedding Given **E**_*i*_ ∈ ℝ^*M*^, we construct

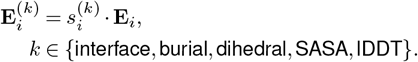

where 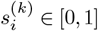 is the biophysical score for metric *k*. The set 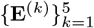 constitutes the enriched embedding space used in the downstream tasksDing et al. (2021). As illustrated in Figure 1, each scaled embedding is projected into a lower-dimensional space, regularized through dropout and non-linear activation. A cross-embedding multihead attention mechanism then integrates the five streams, allowing the model to reweight and combine information across biophysical perspectives.Ding et al. (2021); Xu et al. (2019) Finally, an attention pooling layer aggregates these signals into a compact representation. This design introduces minimal parameters, making it well-suited for limited datasets such as SKEMPI2.

To further guide learning, we impose a multi-task weighted loss that conditions the network to recover not only experimental ΔΔ*G* values but also the interface structural consistency metric ilDDT Mariani et al. (2013); Thomas et al. (2025). This auxiliary prediction encourages the extraction of structurally meaningful features and improves generalization.

Overall, ProtBFF can be applied as a drop-in replacement for the feed-forward modules placed after protein embeddings in ΔΔ*G* prediction pipelines. Because it requires only minimal adaptation to embedding dimensionality, it can readily be integrated with a wide range of pretrained encoders and extended to other downstream structural prediction tasks, offering a general and practical means of injecting biophysical inductive bias into deep learning models.

#### Embedding-aware attention Network Architecture

The ProtBFF framework operates by processing embeddings generated by a pre-trained encoder. For each protein complex, 5 per-residue biophysical scores are computed, each associated with their own scaled embedding. The *k*-th of the five embeddings is denoted by **E**^(*k*)^ ∈ ℝ^*L×M*^, where *L* is the number of residues in the protein complex and *M* is the dimensionality of the embeddings. *L* corresponds to the full sequence length of the complex.

Although the specific embedding dimensionality *M* may differ across different encoders, the embedding-aware attention network is agnostic to this and can be adapted by adjusting its input size hyperparameter. All embeddings are of ℝ^*L×M*^, thereby allowing for per-residue scaling and subsequent pooling operations.

Once the per-score embeddings are obtained, they are max-pooled along the residue dimension, producing five fixed-length vectors in ℝ^*M*^. These are concatenated to form a single representation **X** ∈ ℝ^5*M*^, which is passed into the embedding-aware attention network. The network first reshapes **X** back into its 5 constituent ℝ^*M*^ embeddings, which then each undergo an attention pooling operation. The resulting 5 pooled vectors are processed via a cross-embedding attention mechanism, which attends over all embeddings simultaneously, and normalized to produce a single ℝ^512^ representation. This final representation is passed through a three-layer multilayer perceptron (MLP) to produce ΔΔ*G* predictions. The reported value is the average of the predictions for the forward and reverse mutations to preserve anti-symmetry:

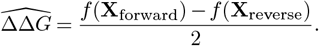

For a full schematic of the ProtBFF framework, please refer to Figure 1

It is possible to fine-tune the model architecture based on the amount of data or the specific task, while maintaining the general methodology—namely, the integration of biophysical features followed by attention-pooling.

### Definition of Biophysical Scores

We define five biophysical scores, which are generated from the wildtype and mutant structures, as follows:

The ***interface score*** is a normalized metric that quantifies how close a given residue is to a protein-protein interface within a protein complex. A higher interface score indicates that the residue is closer to the protein-protein interface and plays a more significant role in inter-chain interactions, and, therefore, a mutation at that residue would be more influential on structural stability Zhao et al. (2025); Mirabello and Wallner (2024).

The interface score for residue *i* is

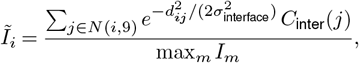

where *N* (*i*, 9) are the 9 nearest neighbors (including *i*), *d*_*ij*_ is the residue–residue distance, and *σ*_interface_ controls the Gaussian width. The inter-chain contact count is

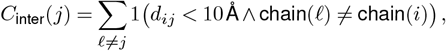

and max_*m*_ *I*_*m*_ is the maximum raw score (the numerator) across all residues. This normalization yields values between 0 and 1 for direct comparison across residues.

The ***burial score*** quantifies how deeply buried a given residue is within a protein, informed by the intuition that mutations to more buried residues generally cause more significant conformational changes Yu et al. (2024). A higher burial score indicates a residue is more embedded in the protein’s interior.

The burial score for residue *i* is

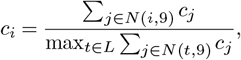

where *N* (*i*, 9) are its 9 nearest neighbors and *c*_*j*_ is the count of residues within 10 Å of residue *j*. Normalizing by the maximum neighbor-sum over all residues *L* yields values between 0 and 1.

The ***dihedral score*** quantifies structural changes in side-chain dihedral (chi) angles after mutation. Mutations causing significant chi angle alterations can lead to substantial conformational rearrangements, impacting protein stability and function. A higher dihedral score indicates a more pronounced structural shift.

The dihedral score for residue *i* is computed by comparing side-chain dihedral angles between the wildtype and optimized structures. Specifically, for each residue *j* ≠ *i* and each of its side-chain dihedral angles *χ*_*k*_, we compute the angular deviation as

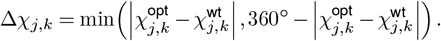

where 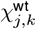 and 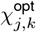 denote the *k*-th side-chain dihedral angle (*χ*) of residue *j* in the wildtype and optimized structures, respectively. This definition selects the minimum angular distance while accounting for angular periodicity. Symmetric 180^°^ flips in aromatic residues (PHE, TYR, HIS) are treated as equivalent conformations and are therefore ignored.

The total angular change associated with residue *i* is obtained by summing over all other residues:

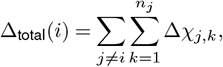

where *n*_*j*_ is the number of side-chain dihedral angles for residue *j*. The dihedral score is then normalized by the maximum total angular change across all residues:

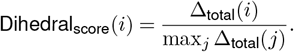

The SASA score quantifies a residue’s exposure to solvent in a protein structure, emphasizing surface accessibility rather than interior packing—with higher values indicating greater exposure Gao et al. (2021). This score is calculated using the *freesasa* package, which determines the Solvent Accessible Surface Area of each residue based on atomic coordinates in the wild-type structure Mitternacht (2016). The SASA score incorporates both the solvent exposure of the residue of interest and the exposure of its nearby residues, weighted by their sequential proximity.

The SASA score for residue *i* is the Gaussian-weighted sum of solvent-accessible surface areas for its 10 nearest sequential neighbors, normalized to [0, 1]:

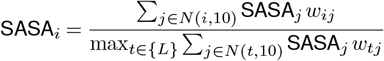

Here, SASA_*j*_ is the solvent-accessible surface area (from *freesasa*) of neighbor *j*, 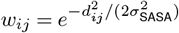 is a Gaussian weight favoring closer neighbors, *N* (*i*, 10) is the 10 nearest sequential neighbors to the left and right of *i*, and *{L}* is the set of all residues.

The ***lDDT score*** measures per-residue atomic conformational changes between the wildtype (template) and FoldX-generated mutant (predicted) structures Mariani et al. (2013). Larger changes indicate residues more likely to affect overall structure and function. The score is normalized to [0, 1], and we scale embeddings by 1 − lDDT so that residues with greater changes receive higher emphasis during max pooling.

#### Mutant Structures

FoldX was used to generate mutant protein structures for all complexes in the SKEMPI2 dataset. After applying RepairPDB twice to the wildtype structures, FoldX relaxes the mutant side chains using the BuildModel function Schymkowitz et al. (2005). Importantly, this procedure does not modify the protein backbone. Although not explored here, alternative methods that incorporate backbone relaxation, such as Rosetta Relax, may produce more accurate mutant structures (albeit at higher computational cost) which could in turn yield improved biophysical features and enhance ΔΔ*G* prediction accuracy Alford et al. (2017).

#### Attention-model Training and Evaluation

The embedding-aware attention network is trained on wildtype and mutant embeddings and includes two MLP heads: one predicting ΔΔ*G* and one predicting interfacial lDDT (ilDDT), a scalar structural consistency metric restricted to interchain contacts Mariani et al. (2013); Biasini et al. (2013). The ilDDT head serves as a regularizer, encouraging the model to extract structurally meaningful features alongside binding energetic signals. Joint training is implemented using a weighted mean squared error loss:

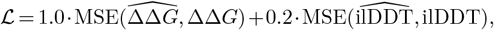

where 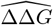 and 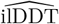 are the network predictions, and the ground-truth ilDDT is calculated by comparing the FoldX-mutated structure to the wildtype PDB template. The use of two output heads ensures that conditioning occurs early in the network, at the cross-embedding attention stage, after which the ΔΔ*G* head can specialize in energetic predictions Girdhar and Ramanan (2017); Lee et al. (2019). During inference, the ilDDT output is ignored, as it serves only as a conditioning signal during training.

### ΔΔ*G* Predictor Benchmarking

We benchmarked eight baseline models, several of which are regarded as state-of-the-art, and evaluated all models using ten-fold cross-validation. For ESM3, both sequence and structures were embedded Hayes et al. (2024), while for ESM2, only sequences were embedded Lin et al. (2023). Although ProMIM and DDAffinity incorporate features and code from RDE-Network, their architectures differ substantially and have been reported to outperform it, making the comparison particularly relevant Mo et al. (2024); Luo et al. (2023); Yu et al. (2024).

#### Model Training and Testing

Benchmarking for consistent comparison across all models was performed on the 60% sequence similarity split of the SKEMPI2 dataset. The models were benchmarked on the entire dataset, including both single and multiple point mutations.

Models not requiring retraining on SKEMPI2 (namely ESM2, ESM3, and ProSST) were used out-of-the-box, while those requiring retraining (ProMIM, RDE-Network, RDE-Linear and DDAffinity) were retrained across all ten folds according to their respective protocols. When ProSST, ESM2, and ESM3 were adapted to the ProtBFF framework, the encoders were not retrained; instead, their embeddings were used to train the attention network for ΔΔ*G* prediction.

For the general benchmarking, the ESM2 model used was esm2_t33_650M_UR50D. For the specific benchmarking across ESM2 model sizes, the models evaluated were esm2_t30_150M_UR50D, esm2_t33_650M_UR50D, esm2_t36_3B_UR50D, and esm2_t48_15B_UR50D.

#### Ablation Study on Biophysical Feature Contributions

To evaluate the relative contributions of the five biophysical features used in ProtBFF, we performed a systematic ablation study. Each variant of the model was trained with one or more features removed at a time, while keeping the remaining components intact. This allowed for the quantification of their individual and collective importance in driving ΔΔ*G* prediction performance.

We used the ProSST backbone as the base model for this analysis and trained each ablated ProtBFF variant on the SKEMPI2 dataset clustered at 60% sequence identity, following the same 10-fold cross-validation protocol described in section*.

#### Out-Of-Distribution Evaluation

For further benchmarking, we used the DMS dataset from Bloom paper, which quantifies changes in binding affinity between the SARS-CoV-2 RBD, the mutational target, and three binding partners: the human ACE2 receptor, and the neutralizing antibodies LY-CoV555 and REGN10987 Starr et al. (2020). The corresponding template structures used for these complexes were PDB IDs 7W9I (RBD–ACE2), 7KMG (RBD–LY-CoV555), and 9LYP (RBD–REGN10987). Although additional antibodies are present in the dataset, they were excluded because their complex sizes exceed the context window limits of ESM and ProSST, or because their corresponding template structures are sparsely represented in the PDB, preventing reliable generation of structural embeddings. After removing entries with missing measurements and filtering out mutations located in structurally unresolved regions, the final datasets comprised 3,669 datapoints for the RBD–ACE2 interaction, 3,156 datapoints for RBD–REGN10987, and 3,137 datapoints for RBD–LY-CoV555. The aforementioned PDB structures were used as wildtype templates, and all mutations were introduced using FoldX following the same protocol applied to SKEMPI2.

To benchmark performance on this dataset, we loaded the models trained during the SKEMPI2 benchmarking. Specifically, models from the first fold of the ten-fold cross-validation were selected for consistency. These pretrained models were then progressively fine-tuned on the Bloom DMS datasets using increasing training set sizes, following the protocol described earlier and illustrated in Figure 3.

