## Supplementary Information for "A General Framework for Injecting Biophysical Priors into Protein Embeddings"

#### This PDF file includes:

Supporting text
Figs. S1 to S2
Table S1
SI References

### Supporting Information Text

**Rationale for Biophysical Score Selection.** To identify which structural and biophysical properties most strongly correlate with binding affinity changes, we systematically evaluated a comprehensive set of scalar features computed at the protein-complex level. All features were calculated by comparing FoldX-generated mutant structures to the experimental wildtype template structures from the Protein Data Bank. Each feature was tested for its correlation with experimental $\Delta\Delta G$ values across the SKEMPI2 dataset, with outliers removed to ensure robust statistical estimates. Table S1 summarizes the Spearman correlation coefficients and associated p-values for all tested features.

Several features demonstrated statistically significant correlations with $\Delta\Delta G$. The interface local distance difference test (ilDDT) exhibited the strongest correlation, followed closely by several other scores. While these correlations are modest in absolute magnitude, they indicate that structural features contain genuine signal about mutational effects on binding affinity.

Importantly, these correlations reflect protein-level scalar summaries. We hypothesized that substantially more information could be extracted by computing features at the per-residue level, enabling the model to selectively weight residues based on their local structural context. This observation motivated our design choice to scale per-residue embeddings rather than simply appending global features to learned representations.

We also evaluated AlphaFold 3–derived predictive metrics, predicted TM-score (pTM), interface predicted TM-score (ipTM), and ranking confidence scores, to assess whether structure prediction confidence measures could provide informative priors for $\Delta\Delta G$ prediction. However, these metrics showed weak or negligible correlations with experimental binding affinity changes. This result is consistent with prior findings that AlphaFold models, while highly accurate for wildtype structure prediction, do not reliably predict the energetic consequences of point mutations (1, 2). In contrast, template-based metrics such as lDDT, which compare FoldX-generated mutant structures to experimental wildtype templates, proved substantially more informative.

Additional features were tested, including a flexibility score that quantified local backbone mobility around the mutation site, calculated by *Flexpert* (3) and a shift score that measured the changes in residue contacts (see Figure S2). However, these exhibited weak correlations with experimental $\Delta\Delta G$ values and did not improve model performance when integrated into ProtBFF. These features were therefore excluded from the final framework. As noted in the main text, this framework can be extended to other relevant biophysical features, which may be specific to tasks outside of $\Delta\Delta G$ prediction.

Based on this analysis, we selected five features for integration into ProtBFF: interface score, burial score, dihedral score, solvent-accessible surface area (SASA), and lDDT. These features were chosen because they (1) capture complementary aspects of protein structure and energetics, and (2) can be computed efficiently at the per-residue level, enabling fine-grained modulation of learned embeddings, and (3) substantively improved the accuracy of the $\Delta\Delta G$ predictions made from a given encoder's embeddings, as noted in the ablation studies in the main text.

| Feature | Spearman $\rho$ | p-value |
|---|---|---|
| ilDDT (4) | $-0.343$ | $1.02 \times 10^{-163}$ |
| ICS (5) | $-0.285$ | $2.87 \times 10^{-110}$ |
| DockQ (6) | $-0.284$ | $4.51 \times 10^{-110}$ |
| Interface Similarity (7) | $-0.233$ | $1.33 \times 10^{-75}$ |
| lDDT (4) | $-0.223$ | $2.23 \times 10^{-67}$ |
| IPS (5) | $-0.177$ | $1.10 \times 10^{-41}$ |
| Ranking Score (1, 8) | $-0.099$ | $3.98 \times 10^{-14}$ |
| pTM (1, 8) | $+0.031$ | $1.76 \times 10^{-2}$ |
| ipTM (1, 8) | $+0.001$ | $9.19 \times 10^{-1}$ |

**Table S1. Spearman correlation coefficients between scalar biophysical features and experimental $\Delta\Delta G$ values on SKEMPI2 (outliers removed). All structural features were computed by comparing FoldX-generated mutant structures to experimental wildtype template structures. Features are ranked by absolute correlation magnitude. Template-based structural metrics (ie. ilDDT, ICS, and DockQ) exhibit the strongest correlations, while AlphaFold 3 prediction confidence metrics (ipTM, pTM) show negligible associations with binding affinity changes.**

**Integration Strategy: Embedding Scaling vs. Alternative Approaches.** Having identified informative biophysical features, we explored multiple strategies for integrating them into deep learning models. The central challenge was to combine pretrained residue-level embeddings, which encode rich sequential and structural information, with explicit biophysical priors in a manner that preserves the strengths of both representations while avoiding information loss or overfitting.

We tested several alternative integration methods prior to arriving at the embedding-scaling approach used in ProtBFF. Feature concatenation, in which biophysical feature vectors were appended directly to residue embeddings either as additional dimensions or as separate global descriptors, failed to provide consistent improvements and in some cases degraded performance. We posit that concatenation treats learned embeddings and biophysical features as independent information channels, preventing the model from using structural priors to selectively emphasize or de-emphasize specific residues. Additive feature integration, in which scalar biophysical values were added as auxiliary input vectors either before or after embedding extraction, similarly underperformed, likely because additive combinations do not inherently modulate the relative importance of residues based on their structural context.

Non-attention architectures, specifically simpler feed-forward multilayer perceptron (MLP) architectures that processed concatenated embeddings and features through fully connected layers, consistently underperformed attention-based variants. The poor performance of purely feed-forward models likely reflects the complex, non-linear interactions between biophysical features and their context-dependent effects on binding affinity (9). Classical machine learning methods, including gradient-boosted trees and support vector regression using biophysical features as direct inputs without pretrained embeddings, performed extremely poorly (Spearman $\rho < 0.20$), underscoring the limitations of relying solely on hand-crafted features without the rich representational capacity of deep embeddings (10). This result is consistent with the known non-linear, high-dimensional nature of protein energetics, which classical methods struggle to capture effectively (11, 12).

Ultimately, we found that multiplicative scaling of embeddings by biophysical scores provided the most effective integration strategy. By scaling each residue's embedding according to its structural relevance—whether measured by interface proximity, burial depth, conformational change, or solvent exposure—the model is guided to focus on residues most likely to influence binding affinity. This approach preserves the full information content of pretrained embeddings while introducing an interpretable structural bias that aligns with physical intuition.

The use of cross-embedding attention further enhances this strategy by enabling the model to construct mixed representations that capture non-linear interactions among biophysical features (13). For instance, a mutation at a buried interface residue may have compounding effects that cannot be captured by considering burial and interface proximity independently. The attention mechanism learns to weight and combine these signals adaptively, producing richer and more context-sensitive residue representations than any single feature or simple feature combination could provide (14).

Together, the embedding-scaling and attention-based integration framework implemented in ProtBFF reflects a principled synthesis of learned representations and physically grounded priors, enabling models to generalize more effectively while remaining interpretable and computationally efficient.

**Extensibility and Adaptability of the ProtBFF Framework.** A key advantage of the ProtBFF framework is its modular design, which enables straightforward adaptation to diverse protein analysis tasks and incorporation of alternative biophysical features or encoders. The framework's three core components—biophysical feature selection, encoder choice, and attention-based integration—are designed to be independently tunable, providing flexibility for future applications.

The five biophysical scores $\{s_i^{(k)}\}_{k=1}^5$ used in this study (interface score, burial score, dihedral score, SASA, and lDDT) were selected based on their empirical correlations with $\Delta\Delta G$ and computational accessibility. However, the framework imposes no constraints on the number or identity of features. For a generalized set of $K$ biophysical scores, the scaled embeddings are computed as

$$\mathbf{E}_i^{(k)} = s_i^{(k)} \cdot \mathbf{E}_i, \quad k \in \{1, 2, \ldots, K\},$$

where $\mathbf{E}_i \in \mathbb{R}^M$ is the residue embedding and $s_i^{(k)} \in [0, 1]$ is the $k$-th biophysical score for residue $i$. Alternative or additional structural descriptors—such as electrostatic potential, hydrophobic moment, secondary structure propensity, or evolutionary conservation scores—can be readily substituted or added without modifying the core architecture. The cross-embedding attention mechanism dynamically learns to weight and combine whichever features are provided, making the framework agnostic to the specific choice of biophysical priors. This modularity enables practitioners to tailor feature selection to domain-specific knowledge or dataset characteristics.

ProtBFF operates on residue-level embeddings and is therefore compatible with any encoder $f_\theta : \mathcal{S} \to \mathbb{R}^{L \times M}$ that maps a protein sequence (or sequence-structure pair) $\mathcal{S}$ to a matrix of per-residue representations, where $L$ is the sequence length and $M$ is the embedding dimension. In this work, we demonstrated integration with ProSST, ESM2, and ESM3, but the framework can equally accommodate other pretrained models such as ProtTrans (15) or future protein language models. The only requirement is that the encoder outputs fixed-dimensional embeddings $\mathbf{E}_i \in \mathbb{R}^M$ for each residue $i \in \{1, \ldots, L\}$. This encoder-agnostic design allows researchers to leverage advances in protein representation learning without redesigning the downstream prediction architecture.

The cross-embedding attention network can be scaled to accommodate tasks with different data availability or complexity. For problems with limited training data, such as the SKEMPI2 benchmark, we employed a relatively compact architecture to avoid overfitting. However, for larger datasets or tasks requiring integration of many biophysical features, the model can be expanded by increasing the number of attention heads $H$, stacking multiple attention layers, or widening the feed-forward networks. Similarly, if higher-dimensional embeddings are used, the attention mechanism naturally scales to process richer representations, though this may require proportional increases in model capacity and computational resources.

All models reported in this study, including those based on ESM2 and ProSST encoders combined with ProtBFF, were trained on a single NVIDIA A40 GPU (48 GB memory). Training times for the attention network ranged from 15 minutes to an hour depending on encoder size and dataset complexity, making the approach accessible for typical academic or industrial computational environments. Inference is similarly efficient, with $\Delta\Delta G$ predictions computed in milliseconds per mutation once embeddings are generated.

This extensibility makes ProtBFF a general scaffold for protein property prediction tasks beyond $\Delta\Delta G$ estimation. By substituting appropriate biophysical features $\{s_i^{(k)}\}$ and target labels $y$ (e.g., replacing binding affinity changes with folding stability, ligand affinity, or catalytic efficiency), and retraining the attention module, the framework could be adapted to predict protein stability ($\Delta\Delta G_{\mathrm{fold}}$), ligand binding affinity, enzyme catalytic efficiency, or other structure-function relationships. The core principle—guiding learned representations with physically interpretable priors through multiplicative scaling—remains broadly applicable across protein engineering and computational biology.

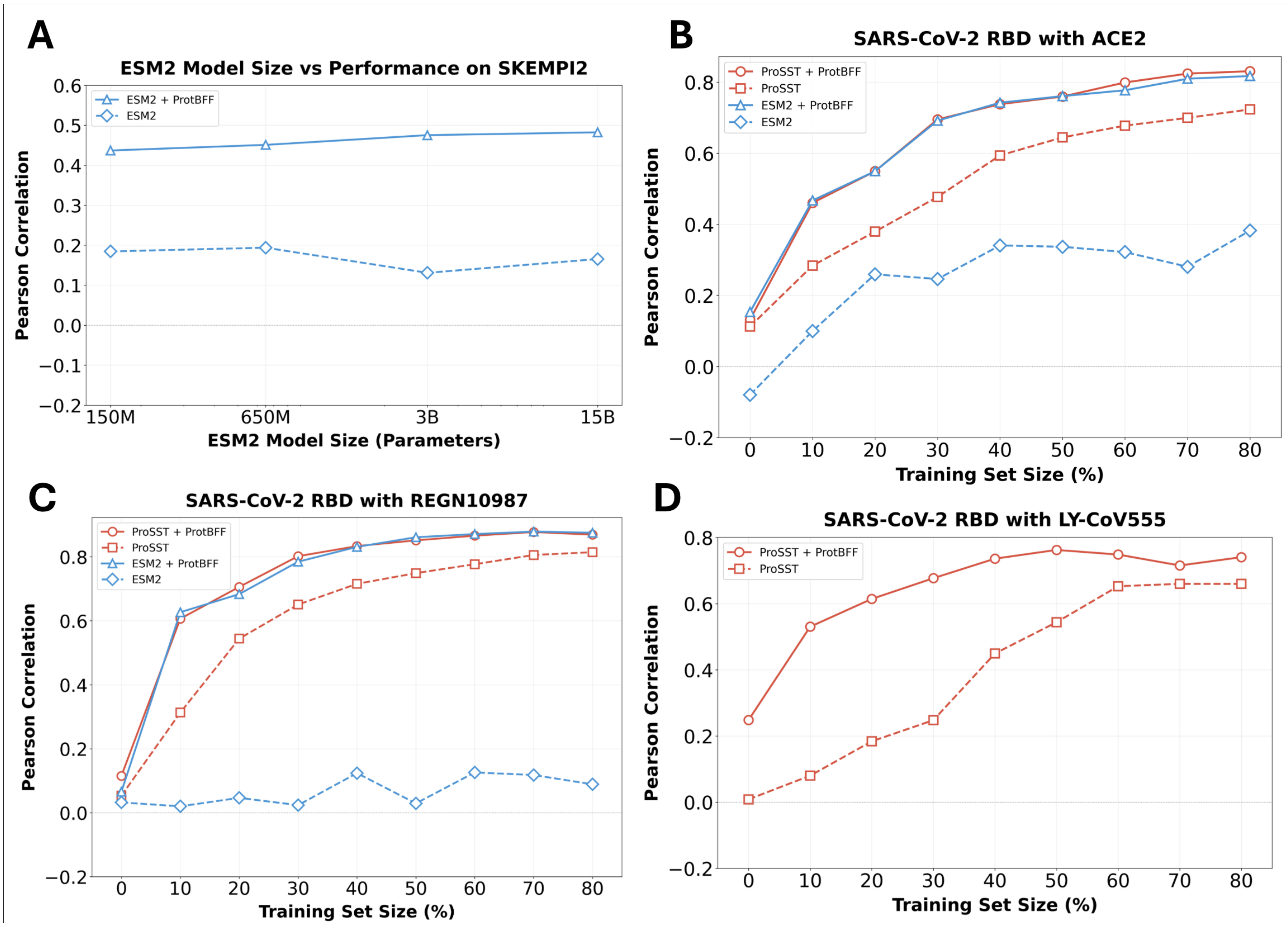


**Fig. S1.** **A** Performance of ESM2 on the SKEMPI2 dataset, measured by Pearson correlation coefficient across model sizes defined by the number of parameters. **B–D** Performance of ProSST and ESM2, measured by Pearson correlation coefficient, with and without ProtBFF, on deep mutational scanning (DMS) datasets of SARS-CoV-2 RBD mutations when bound to **B** ACE2, **C** REGN10987, and **D** LY-CoV555. *Note:* Due to the large size of the RBD–LY-CoV555 complex template structure (PDB 7KMG), it exceeds the context length limit of ESM2 and therefore could not be processed. As a result, ESM2-based results are omitted from subfigure D.

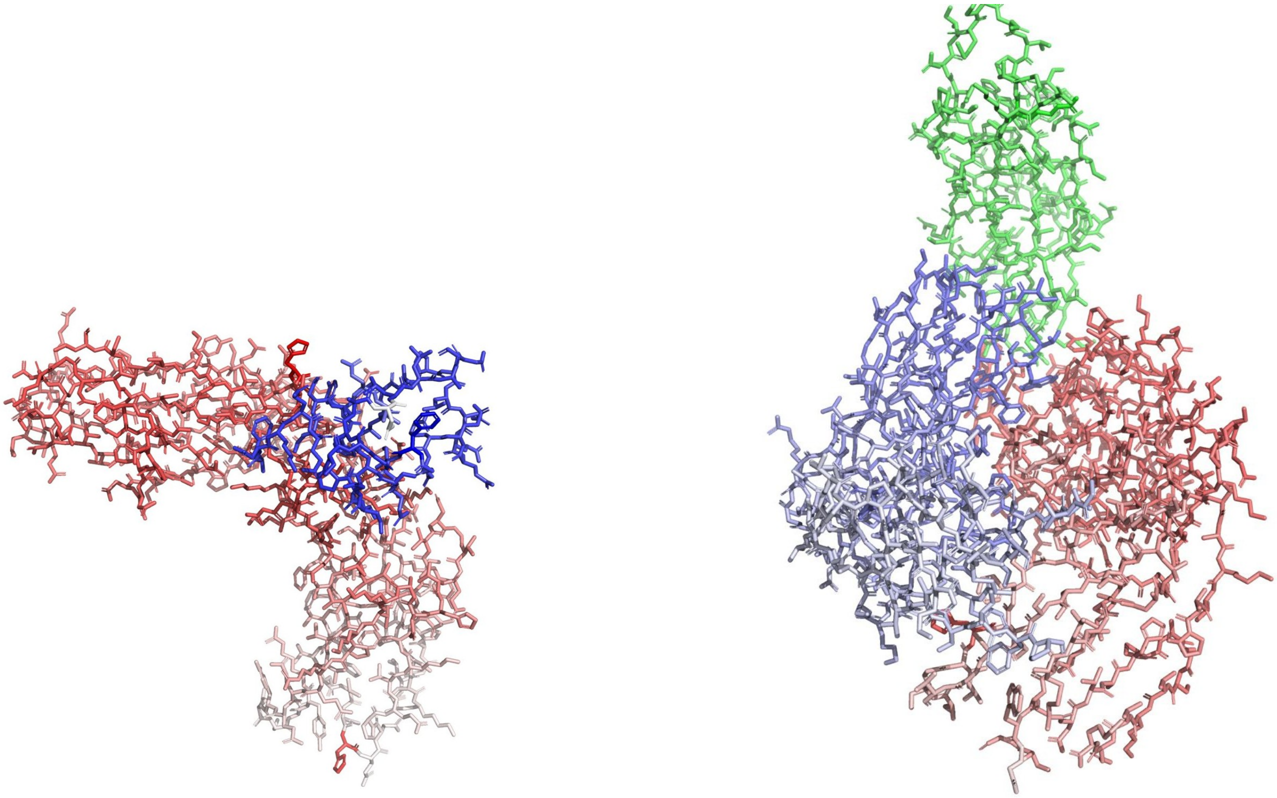

**(a)** 1FC2 Shift Score Representation

**(b)** 1976 Shift Score Representation

**Fig. S2.** Panels A and B display the shift score projected onto the protein structure, with residues colored according to their score: chain A in red, chain B in blue, and chain C (when present) in green. Residues with higher biophysical contributions are shown with more intense colors, while those with lower contributions appear closer to gray. As most residues in both proteins exhibit similar shift scores—reflecting the small perturbations applied by FoldX (16)—the shift score offers limited biophysical discrimination.
